# Improved DOP-PCR (iDOP-PCR): a robust and simple WGA method for efficient amplification of low copy number genomic DNA

**DOI:** 10.1101/128736

**Authors:** Konstantin A. Blagodatskikh, Vladimir M. Kramarov, Ekaterina V. Barsova, Alexey V. Garkovenko, Dmitriy S. Shcherbo, Andrew A. Shelenkov, Vera V. Ustinova, Maria R. Tokarenko, Simon C. Baker, Tatiana V. Kramarova, Konstantin B. Ignatov

## Abstract

Whole-genome amplification (WGA) techniques are used for non-specific amplification of low-copy number DNA, and especially for single-cell genome and transcriptome amplification. There are a number of WGA methods that have been developed over the years. One example is degenerate oligonucleotide-primed PCR (DOP-PCR), which is a very simple, fast and inexpensive WGA technique. Although DOP-PCR has been regarded as one of the pioneering methods for WGA, it only provides low genome coverage and a high allele dropout rate when compared to more modern techniques. Here we describe an improved DOP-PCR (iDOP-PCR). We have modified the classic DOP-PCR by using a new thermostable DNA polymerase (SD polymerase) with a strong strand-displacement activity and by adjustments in primers design. We compared iDOP-PCR, classic DOP-PCR and the well-established PicoPlex technique for whole genome amplification of both high- and low-copy number human genomic DNA. The amplified DNA libraries were evaluated by analysis of short tandem repeat genotypes and NGS data. In summary, iDOP-PCR provided a better quality of the amplified DNA libraries compared to the other WGA methods tested, especially when low amounts of genomic DNA were used as an input material.

## Introduction

Molecular analysis of limited quantities of genomic DNA (gDNA) is crucial for characterization of single cell genomes, in preimplantation genetic diagnosis (PGD), in DNA forensics and many other applications.

Genomic DNA can be analyzed by a variety of methods: next-generation-sequencing (NGS), microarrays, multiplex STR (short tandem repeat) genotyping, or parallel qPCR techniques addressing multiple genomic regions. However, these analyses require a small but significant amount of human gDNA, in the range of 1 to 100 ng. This corresponds to 160–16000 human cells [1] and so these approaches are not appropriate to the analysis of single-cell genomes.

For samples with limited DNA content, a step of DNA amplification could be used to facilitate further analysis. Whole genome amplification (WGA) is an *in vitro* method to amplify gDNA and is thus useful in order to obtain sufficient material for analyses of low copy number gDNA (<100 pg), the range typically found when isolating DNA from single cells [1]. The current WGA techniques involve one of two approaches: isothermal amplification of DNA or thermo-cycling (PCR-based) methods. Detailed descriptions and comparisons of the different WGA methods can be found in many reviews [2-5] and research articles [6-12].

Multiple displacement amplification (MDA) is the main method for isothermal WGA. This method uses random hexamer primers and bacteriophage Phi29 DNA polymerase, which exhibits strong DNA displacement capabilities [6].

The main techniques used in PCR-based methods are degenerate oligonucleotide-primed PCR (DOP-PCR) [7], multiple annealing and looping based amplification cycles (MALBAC) [8] and the PicoPlex technique [9]. The principle of DOP-PCR is to use a single primer containing a central random sequence. DOP-PCR begins with a few pre-amplification cycles at a low initial annealing temperature, facilitating random primer annealing. Pre-amplification is then followed by PCR amplification of these initial DNA fragments. Currently, the best-in-class performance for PCR-based WGA methods is achieved with MALBAC and PicoPlex techniques. Both methods are very similar [2, 3] and, in contrast to DOP-PCR, utilize different kinds of primers/enzymes for a pre-amplification of DNA (the library generation step) and for PCR-amplification of the DNA fragments generated (the library amplification step) [8, 9]. The two-step protocols of MALBAC and PicoPlex are more labor-intensive than the DOP-PCR procedure, but provide much superior WGA performance when characteristics such as allele drop out rate and genome coverage [3] are considered.

In earlier work we reported improvements in DNA amplification by the use of SD DNA polymerase, a thermostable DNA polymerase with a strong strand-displacement activity [13]. Here, we describe a new variant of DOP-PCR with enhanced WGA performance, which is achieved by SD polymerase application. We also compare improved DOP-PCR (iDOP-PCR) with “classic” DOP-PCR [7] and with a commercially available PicoPlex technique from Rubicon Genomics Inc. (MI, USA), which is currently the predominant method used for preimplantation genetic diagnosis (PGD) and other medical applications [14, 15].

## Materials and methods

### Enzymes and reagents

SD DNA polymerase, Taq DNA polymerase and the reaction buffers were supplied by Bioron GmbH, Ludwigshafen, Germany (www.bioron.net). dNTPs were obtained from Bioline Limited (London, GB).

The PicoPLEX WGA Kit, developed and manufactured by Rubicon Genomics, Inc., was supplied by New England Biolabs, Inc. (Ipswich, MA, USA).

The COrDIS Plus STR Amplification Kit was obtained from *Gordiz LLC* (Moscow, Russia, http://gordiz.ru/index.php/en/).

Oligonucleotide primers for DOP-PCR and iDOP-PCR were synthesized by *Evrogen JSC* (Moscow, Russia).

Human gDNA (obtained from one individual) was supplied by *Syntol JSC* (Moscow, Russia). Nobody working on this project was included as a sample donor for any experiments described herein. The concentration of the human gDNA was verified using the Quant-iT™ PicoGreen® dsDNA Assay Kit (Molecular Probes, Inc., Eugene, OR, USA) and the Applied Biosystems Quantifiler® Human DNA Quantification Kit (Applied Biosystems, Foster City, CA, USA) according to the manufacturers’ instructions.

### WGA-libraries preparation

WGA libraries of human gDNA were prepared by three different methods: PicoPlex, DOP-PCR and iDOP-PCR. Input template gDNA was: 15 ng, 1.5 ng, 0.15 ng, 0.015 ng, and a no template as a negative control. For each sample of input DNA, six separate WGA reactions were performed and six separate WGA libraries were obtained using the methods described above. After the amplifications, the yields of WGA reactions were quantified using the PicoGreen® dsDNA Assay Kit (Molecular Probes, Eugene, OR, USA) and by Agilent 2200 TapeStation Instrument with Genomic DNA ScreenTape System (Agilent Technologies, Waldbronn, Germany).

### PicoPlex Amplification

PicoPlex WGA reactions were carried out using the PicoPLEX WGA Kit (New England Biolabs, Inc., Ipswich, MA, USA). Briefly, gDNA or ddH2O (as a negative control) were added to the Sample Preparation Cocktail and incubated as required by the manufacturer. Pre-amplification was carried out in 15 μl of the Pre-Amp reaction mixture for 12 cycles of: 95°C for 15 sec; 15°C for 50 sec; 25°C for 40 sec; 35°C for 30 sec; 65°C for 40 sec; 75°C – 40 sec. After the pre-amplification stage, 60 μl of freshly prepared Amplification Cocktail was mixed with 15 μl of pre-amplification product. The amplification stage was carried out for 14 cycles of: 95°C for 15 sec; 65°C for 1 min; 75°C – 1 min.

### Classic DOP-PCR Amplification

The reaction mixture (25 μl) for each sample contained: 2 μM DOP primer (5’-CCGACTCGAGNNNNNNATGTGG-3’) as described by Telenius et al. [7], 1x PCR buffer for *Taq* polymerase, 3 mM MgCl_2_, 0.25 mM dNTPs (each), 2.5 U *Taq* polymerase, and 5 μl diluted template gDNA or ddH2O for the negative control. The initial pre-amplification parameters were 95°C for 2 minutes, followed by 5 cycles of: 94°C for 1 minute; 30°C for 1 min; ramp at 0.3°C/s to 72°C and finally 72°C for 3 minutes. This was followed by a PCR amplification of 35 cycles of: 94°C for 30 sec; 56°C for 30 sec and 72°C for 2 minutes. The PCR amplification was completed by an incubation at 72°C for 5 minutes.

### iDOP-PCR Amplification

The reaction mixture (25 μl) for iDOP-PCR contained: 0.4 μM iDOP primer (5’-GTGAGTGATGGTAGTGTGGAGNNNNNNATGTGG -3’); 1x buffer for SD polymerase; 3 mM MgCl2; 0.25 mM dNTPs (each); 10 U SD polymerase and 5 μl diluted gDNA (or ddH2O for the negative control). The initial pre-amplification parameters were 92°C for 2 minutes, followed by 6 cycles of: 92°C for 1 minute; 30°C for 1 min; ramp at 0.3°C/s to 68°C; 68°C for 3 minutes. This was followed by a PCR amplification step of 14 cycles of: 92°C for 30 sec; 62°C for 30 sec; 68°C for 3 minutes. The PCR amplification was completed by an incubation of 68°C for 2 minutes.

### Genetic analysis of the obtained WGA libraries by multiplex STR genotyping

Whole genome amplified samples of human DNA and a positive control of the initial, non-amplified human gDNA were analyzed by multiplex STR genotyping. In each sample, 38 alleles (in 19 STR loci and AMEL) were analyzed. Data was verified statistically by analyzing the results of six separate wgaDNA samples (N = 6 × 38 allels = 228 allels) for each starting amount of gDNA amplified by each WGA method obtained from the assay.

The STR genotyping of the samples was performed in *Genetic Expertise LLC* (Moscow, Russia) by COrDIS Plus® STR Amplification Kit according to the manufacturer’s instructions. For the assay, we used 1 ng of each sample’s DNA as a template.

### Genetic analysis of the WGA libraries by next-generation-sequencing (NGS)

Two PicoPlex and two iDOP-PCR WGA products, obtained from 15 pg (about 2.5 genome copies) human gDNA, were selected for further characterization by Next Generation Sequencing (NGS). For this, 500 ng of the WGA products and non-amplified gDNA (as a control sample) were fragmented to an average size distribution of 400 bp with the S220 Focused Ultrasonicator (Covaris Inc., Woburn, MA, USA). Sequencing libraries were generated using NEBNext Ultra™ DNA Library Prep (New England Biolabs, Inc., Ipswich, MA, USA) kits, following the manufacturer’s protocol. The five NGS libraries obtained were quantified with qPCR NEBNext Library Quant Kits (New England Biolabs, Inc., Ipswich, MA, USA) and with the Agilent 2200 TapeStation Instrument with a D1000 Tape System (Agilent Technologies, Waldbronn, Germany). The five libraries were then mixed into one pool. NGS was performed on 4 lanes of an Illumina HiSeq2500® Instrument (Illumina, California, USA) in HighOutput paired-end mode, resulting in 1,189,172,690 reads, each of which was ∼100 nt long. FASTQ files were generated using BCL2FASTQ software v2.17.1.14 (Illumina, California, USA). The FASTQ files were uploaded to NCBI SRArchive under project ID: PRJNA349144.

### Bioinformatic data analysis

The FASTQ files were quality controlled using FASTQC v0.11.4 (Babraham bioinformatics, Cambridge, UK). This revealed a disproportionate oscillation of the percentage of the bases in the first 30 bp of the reads. In addition to trimming these 30 bp, adapters and low quality read ends with Phred quality scores of less than 15 were trimmed with FLEXBAR v.2.5 [16]. Filtered reads with a minimum length of 70 bp were subsequently aligned to the human genome hg19 (with scaffolds removed) using BOWTIE2 software v2.2.6 [17]. Genome coverage and other statistics were calculated using SAMtools v1.0 (http://www.htslib.org/) and BEDtools v2.19.1 (http://bedtools.readthedocs.io) with addition of custom PERL and shell scripts.

Lorenz curves and copy number variation (CNV) detection were performed using the bioinformatics tool GINKGO [18]. For this, the genome sequence was divided into non-overlapping windows or bins of 1 Mb in size and the number of reads per bin was calculated. The number of reads per bin was corrected for the bias introduced by the inability to map reads into repetitive regions of the genome by using mappability tracks created via self-alignment of the reference genome. Copy numbers were estimated under the assumption that for a diploid genome, the majority of the genome has a copy number equal to two. The copy numbers of other regions were then estimated based on the segment ratio relative to two.

## Results and discussion

### Preparation of the WGA libraries

We compared three WGA methods: the “classic” DOP-PCR, a more modern PicoPlex technique and our improved DOP-PCR (iDOP-PCR). For this, we amplified high and low copy number human genomic DNA and analyzed the obtained WGA libraries by STR- and NGS- based assays.

As a template for preparation of WGA libraries, we used multiple series of ten-fold dilutions of human gDNA and added from 15 ng to 15 pg of gDNA per reaction, which corresponded to between 2.5 and 2500 copies of human genome. For each gDNA dilution sample and each method, six separate WGA reactions were performed.

All three methods provided a similar yield of amplified DNA (about 40 - 50 ng/μl) independent of the initial amount of the template (S1 Table).

The WGA libraries were analyzed by agarose-gel electrophoresis (S1 Fig) and by Agilent 2200 TapeStation Instrument (Fig 1). Classic DOP-PCR and PicoPlex provided WGA libraries with a size of DNA fragments in a range 250 – 1500 bp, whereas iDOP-PCR generated amplified DNA with about 4 times larger size distribution of 800 – 6000 bp.

**Fig 1.**
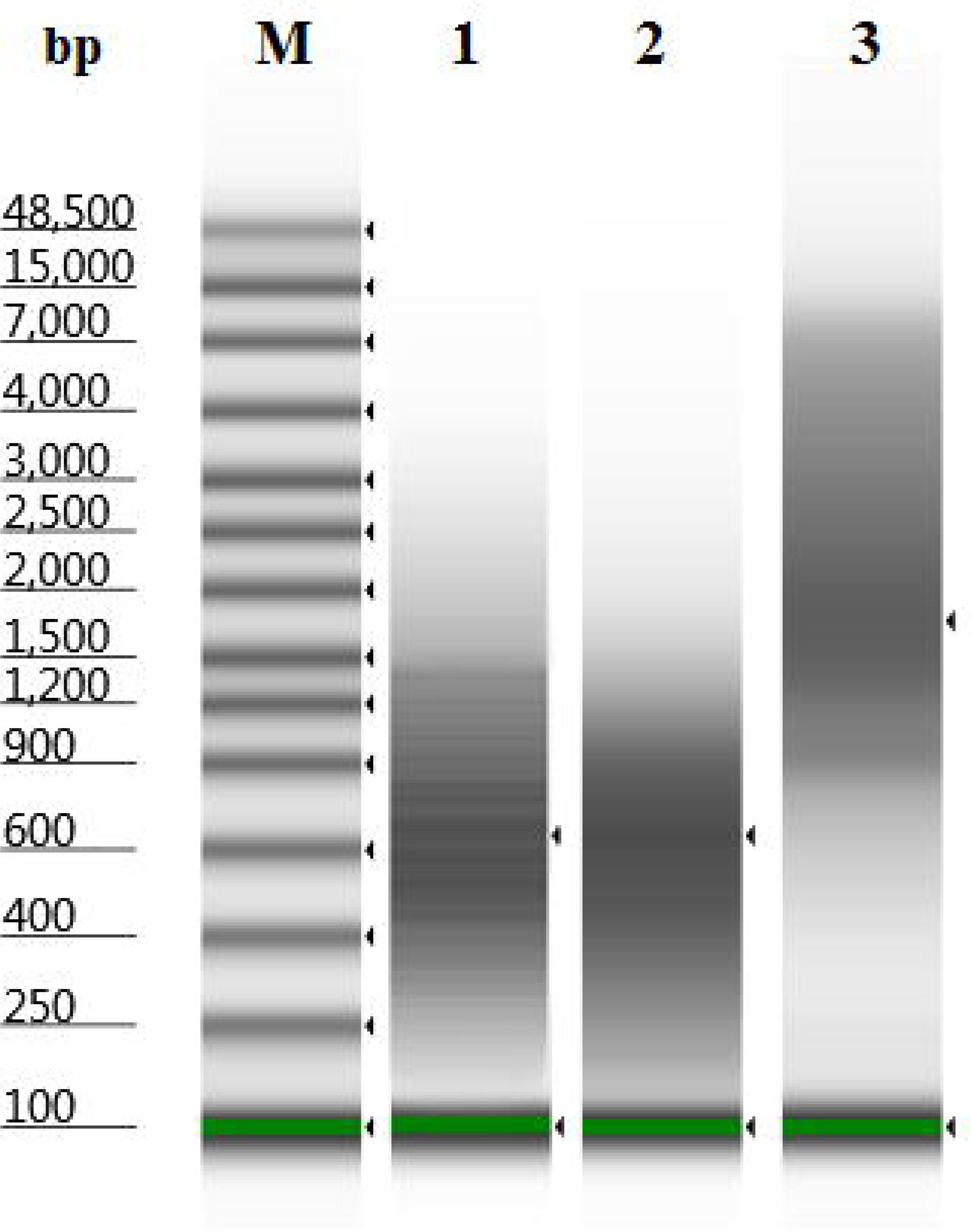
Electrophoretic analysis of WGA libraries by Agilent 2200 TapeStation Instrument. The libraries were obtained by DOP-PCR (lane 1), PicoPlex (lane 2), and iDOP-PCR (lane 3) from 15 pg of human gDNA. **M** – DNA marker “Genomic DNA ScreenTape”.

### STR genotyping assay of human WGA libraries

The important characteristics of WGA in any analysis method (including microarray and PCR based techniques) are allele drop out (ADO) and allele drop in (ADI) rates. ADO and ADI can arise from errors occurring during the amplification. We estimated these characteristics by multiplex STR genotyping of human WGA DNA samples [19], which is a relatively simple and inexpensive method. Non-amplified gDNA was used as a control for the multiplex STR assay. In each DNA sample, 38 alleles (in 19 STR loci and AMEL) were analyzed. Data for each starting amount of gDNA amplified by each WGA method was obtained from the analysis of six separate WGA libraries, giving a total of 228 alleles for statistical calculation of each data point. For estimation of ADO and ADI errors, we calculated drop out for concordant alleles and drop in for discordant alleles. Main results of the assay are summarized in Table 1, detailed data of the assay can be found in S2 Table.

**Table 1.**
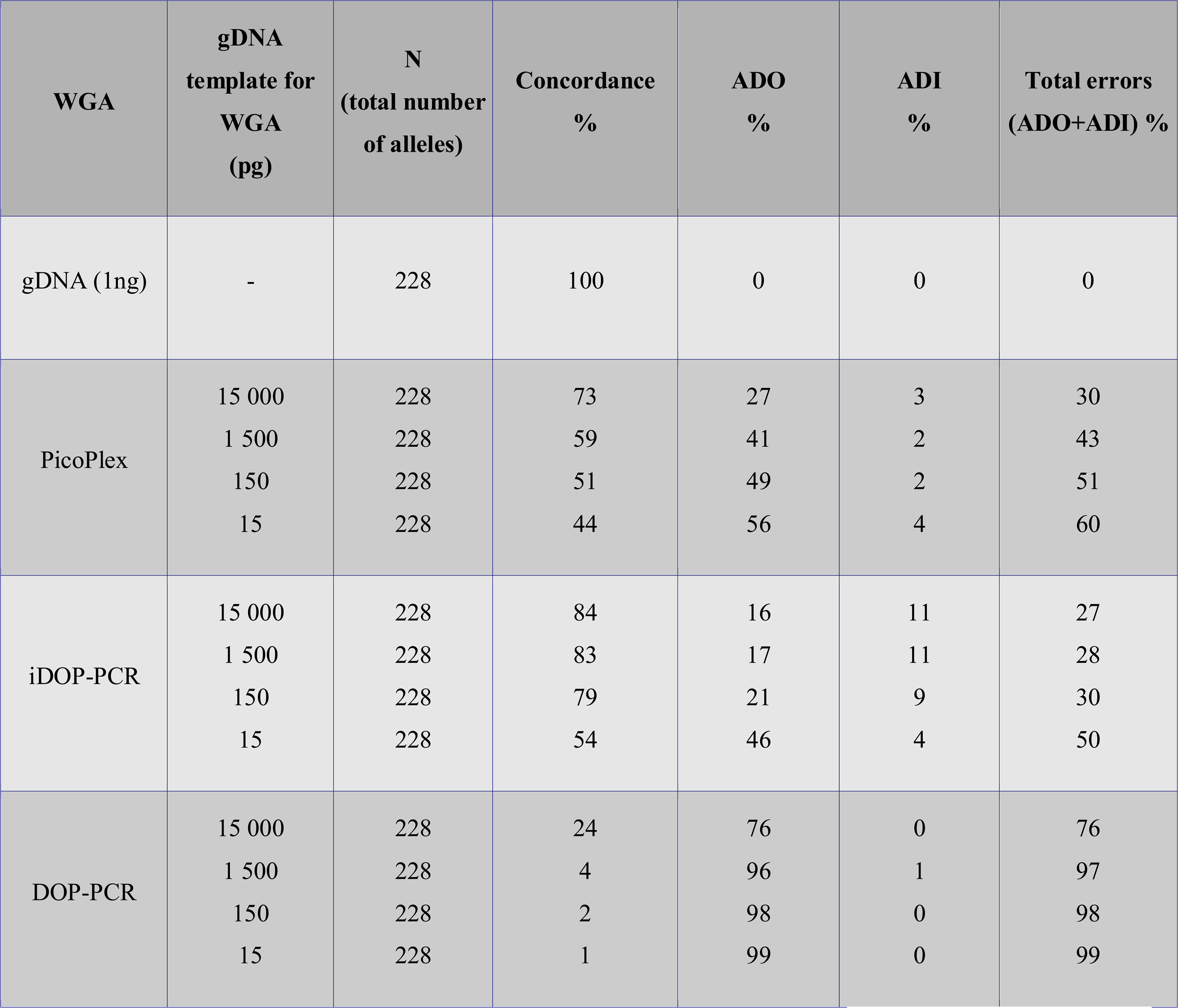
Multiplex STR genotyping of WGA samples and non-amplified gDNA.

In each wgaDNA and non-amplified gDNA sample, 38 allels were analyzed. For each starting amount of gDNA amplified by each WGA method the statistical data was obtained from the assay of six separate wgaDNA samples. Total **N = 6 × 38 allels = 228 allels.** Allele drop out (ADO) and allele drop in (ADI) errors were calculated as drop out for concordant alleles and drop in for discordant alleles.

Among all the WGA methods, classic DOP-PCR exhibited the lowest performance quality in the ADO-ADI test. Amplification of the low amounts of DNA template resulted in very high level of the errors, close to 100% (Table 1), whereas the high copy number template (15 ng gDNA) amplification showed somewhat better concordance rates (24%) between amplified and non-amplified control gDNA. In comparison, PicoPlex generated WGA libraries with better characteristics. High copy number gDNA amplification (from 1.5 – 15 ng gDNA) using the PicoPlex method resulted in 59 – 73% concordance rates between amplified and non-amplified gDNA with a lower rate of error (30 – 43%). Low copy number amplification (from 15 – 150 pg gDNA) resulted in 44-51% concordance rates with non-amplified gDNA and a 51 – 60% error rate.

In these experiments, the modified iDOP-PCR had a better performance than PicoPlex. High copy number template gDNA amplification generated 83 – 84% concordance with non-amplified gDNA and the lowest error rate at 27 – 28%. Low copy number gDNA amplification by iDOP-PCR resulted in 54 – 79% concordance and 30 – 50% error. (Table.1)

Based on the data obtained from the STR assay, further analysis by next generation sequencing (NGS) was restricted to the PicoPlex and iDOP-PCR WGA libraries.

### Comparison of PicoPlex and iDOP-PCR Human WGA libraries by NGS

In addition to allele drop out and drop in rates, other key performance criteria for WGA include reproducibility, genome coverage, uniformity of the amplification and the rate of unmappable sequences. These parameters were compared by NGS analysis of two PicoPlex and two iDOP-PCR WGA libraries obtained from amplification of 15 pg (about 2.5 copies) human gDNA.

To compare the genome coverage of single genome copies with PicoPlex and iDOP-PCR, we used the non-amplified gDNA sequenced at an 8× depth as the reference on the assumption this represented 100% coverage [3]. The comparison was done using the total raw data of 24 Gb for each DNA sample. The raw data generated in this study have been deposited in the National Centre for Biotechnology Information (NCBI) Sequence Read Archive under BioProject accession number PRJNA349144 (https://www.ncbi.nlm.nih.gov/bioproject/PRJNA349144/).

We have shown that the coverage of low copy number genome amplification by PicoPlex WGA was about 51% (which was close to data reported previously [3]) and by iDOP-PCR was about 61% (Table 2).

**Table 2.**
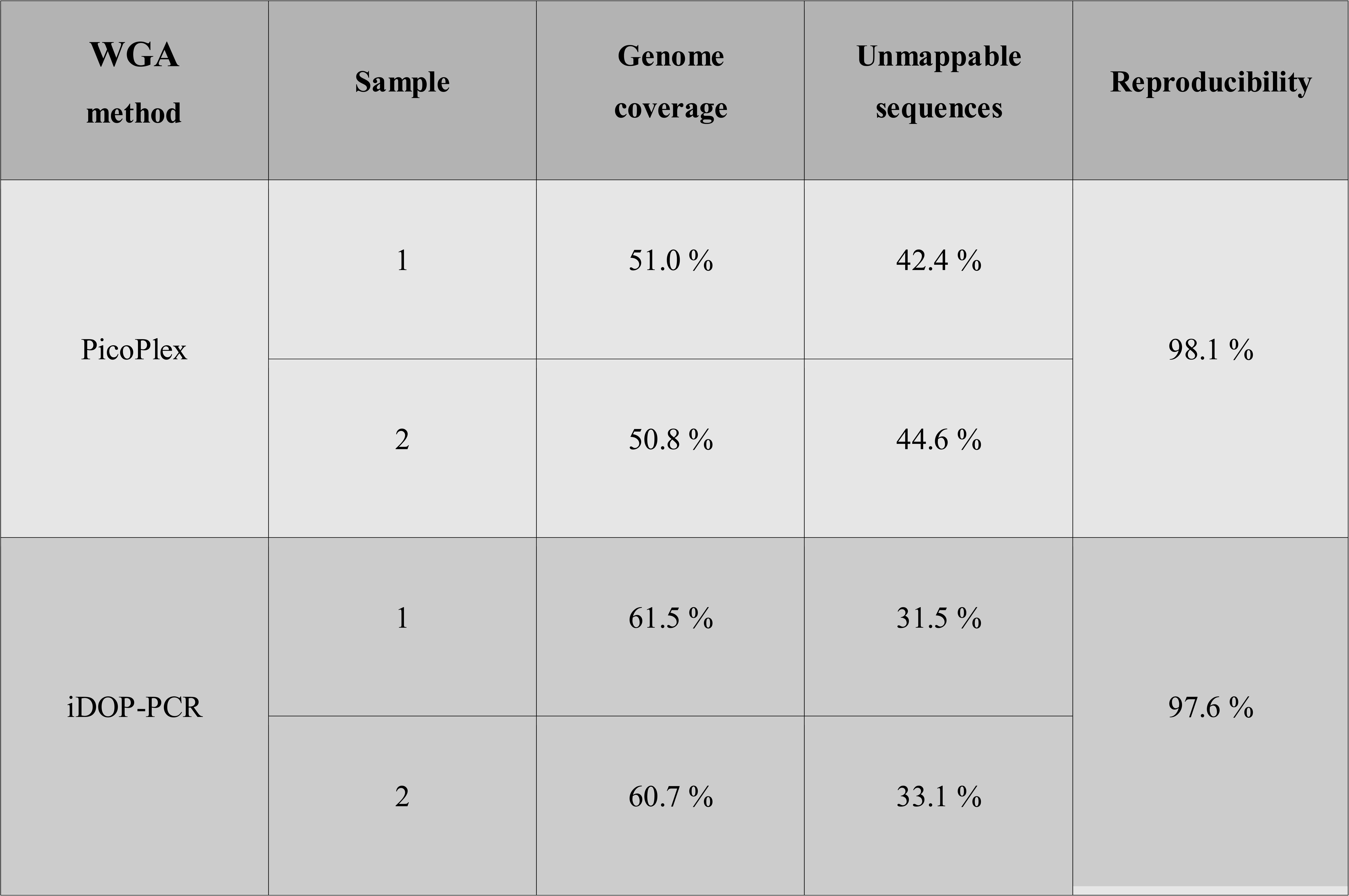
Comparison by NGS the parameters of PicoPlex and iDOP-PCR whole-genome amplification of single-genome-copies.

Key characteristics of the WGA methods (reproducibility of the methods, genome coverage, the rate of unmappable sequences) were compared by NGS analysis of two PicoPlex and two iDOP-PCR WGA libraries. Each library was obtained from 15 pg (about 2.5 copies) Human gDNA.

We found that pooling the raw data from two independently obtained libraries generated by either of the method (PicoPlex or iDOP-PCR) improved the genome coverage by less than 5%. However, when the raw data from libraries generated by two different methods (PicoPlex and iDOP-PCR) was pooled, the genome coverage increased up to 77%. This could indicate that the two WGA methods may differ from each other in which genomic areas they are biased to amplify, at least partially. Thus, the results of NGS could be greatly improved by using a combination of several WGA methods.

As well as coverage, the reproducibility of WGA is a key characteristic in single genome measurements and comparisons. To characterize reproducibility, Pearson’s cross-correlation coefficient of the read densities throughout the genome between two repeated WGA librares was used [3]. This allowed is to show that both PicoPlex and iDOP-PCR provided a high level of reproducibility, at about 98% (Table 2).

Unmappable sequences are generated in the WGA process from the formation of non-template DNA fragments and other nonspecific incorporations/insertions/deletions. A large fraction of unmappable reads reduces the efficiency and the apparent coverage of the genome sequencing. Thus, the unmappable read rate is an important characteristic of the quality of a WGA library. In our experiments, the PicoPlex kit generated 42 – 45% of unmappable sequences, whereas the iDOP-PCR generated only 31 – 33% (Table 2).

Uniformity of the genome amplification is another essential characteristic of WGA methods. Lorenz curves (Fig. 2) were used to evaluate coverage uniformity throughout the genome [8]. The curves give the cumulative fraction of reads as a function of the cumulative fraction of the genome. We compared the Lorenz curves for iDOP-PCR and PicoPlex WGA samples at 8× mean sequencing depth (Figure 2). The curves demonstrated that both iDOP-PCR and PicoPlex methods exhibited very similar uniformity of genome amplification.

**Fig 2.**
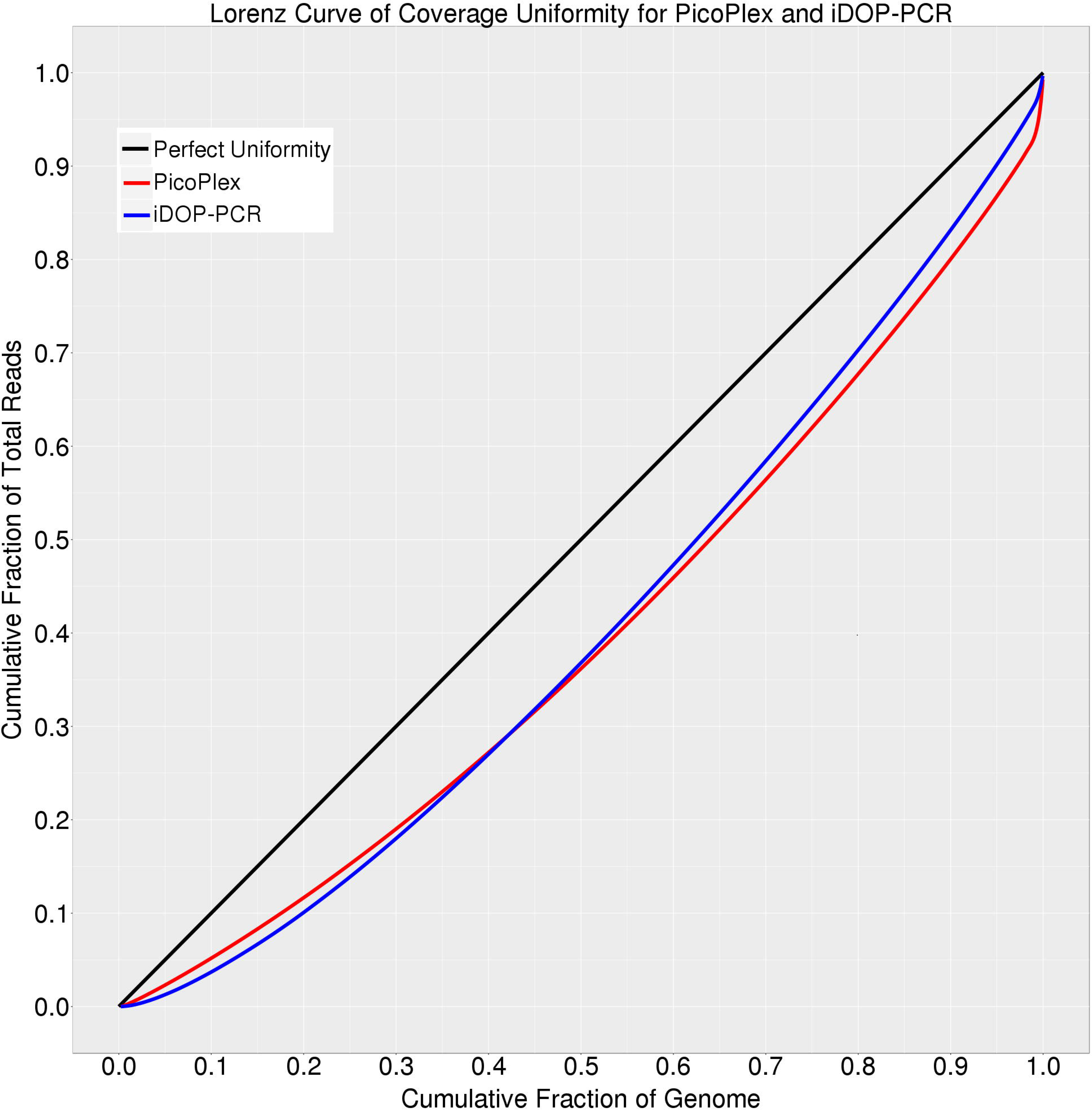
Lorenz curves of PicoPlex and iDOP-PCR WGA samples. A Lorenz curve gives the cumulative fraction of reads as a function of the cumulative fraction of genome. Perfectly uniform coverage would result in a diagonal line (black). PicoPlex (red curve) and iDOP-PCR (blue curve) generate similar deviations from the diagonal as a result of biased coverage. All samples were sequenced at 8x depth.

The ability of WGA to produce samples that are suitable for accurate measurements of copy number variation (CNV) is also very important for further genome evaluation [8, 12]. WGA of low copy number gDNA can lead to a disproportionate amplification of genomic regions. This can result in false positive or false negative copy number changes. We compared the CNV data obtained after iDOP-PCR and PicoPlex amplifications of 15 pg human gDNA. Figure 3 shows the raw read density of all 23 chromosomes and illustrates clearly the sequence-dependent bias along the genome for each WGA sample. Both iDOP-PCR and PicoPlex WGA methods give very similar CNV raw data (Fig. 3).

**Figure 3.**
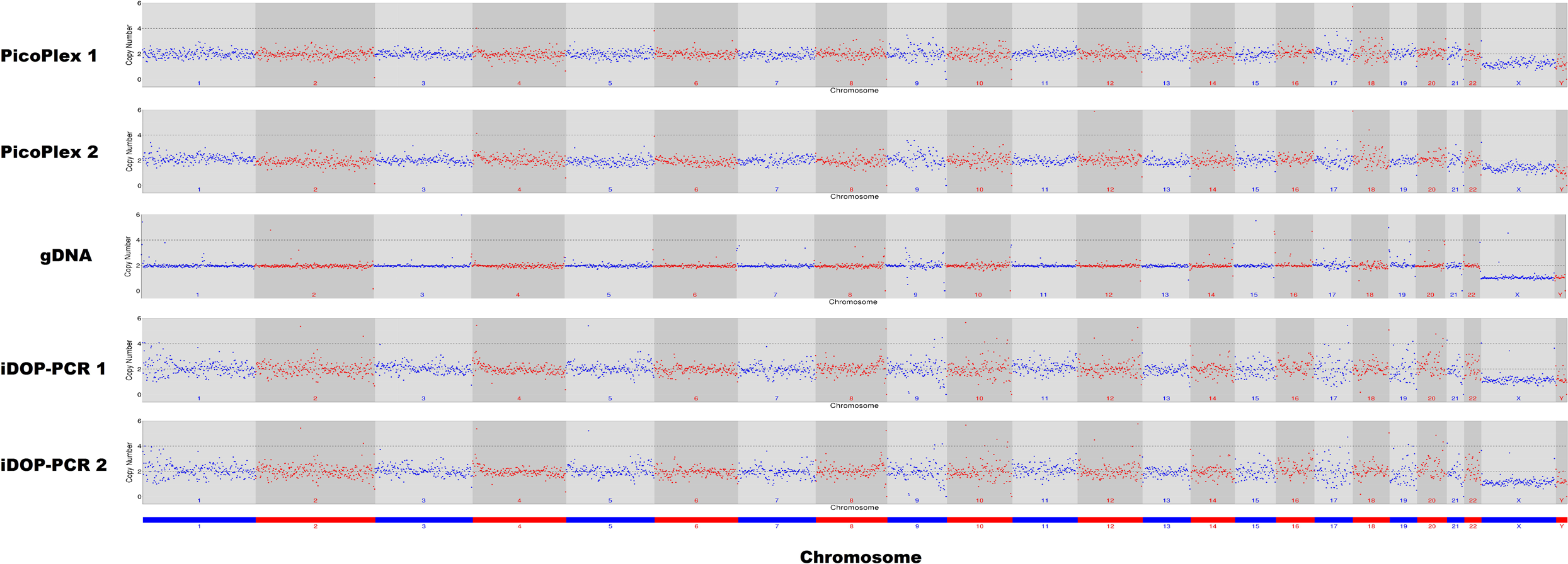
CNVs of diploid human genome from single genome copies amplified by PicoPlex and iDOP-PCR WGA methods. Digitized copy numbers across the genome are plotted for two PicoPlex and two iDOP-PCR WGA samples as well as the non-amplified gDNA sample for control. Raw data at a sequencing depth of 8× with a bin size of 1,000 kb are mapped to the human reference genome. The chromosomes are shown in alternating red and blue colors.

## Conclusions

At first glance, iDOP-PCR looks like a slightly different variant of DOP-PCR method, but the use of DNA polymerase with a strong strand-displacement activity, instead of Taq polymerase, allows to extremely enhance WGA performance of the method. In summary, the comparison of three WGA methods demonstrated clearly that DOP-PCR is unsuitable for WGA of low copy number gDNA (< 1 ng). Both PicoPlex and iDOP-PCR performed well in WGA from single genome copies. Moreover, iDOP-PCR markedly outperformed PicoPlex in the following characteristics: allele dropout (ADO) rate, genome coverage, and amount of unmappable sequences in WGA library, whereas other parameters, such as reproducibility, uniformity and CNV detection were similar for the two methods. It should be noted that at the time of writing, PicoPlex is considered to be theWGA method of choice when ADO, reproducibility, uniformity and CNV detection are of importance [3, 12].

Practically, the greatest advantage of iDOP-PCR lies in its simplicity and cost-effectiveness, being no more complex than ordinary PCR and requiring little investment in kits or reagents. High reproducibility and low ADO rates indicate its potential suitability for some medical applications such as preimplantation genetic diagnosis (PGD). PicoPlex and MALBAC, which are widely used for PGD, utilize two different types of primers and two different enzymes for pre-amplification and amplification stages of WGA. As a result, reaction tubes in these older methods are opened at least twice during WGA, reducing the ease of application to high-throughput analysis and increasing the risk of cross-contamination. In contrast, iDOP-PCR utilizes one primer and one enzyme for all WGA stages and does not require multiple manipulations during WGA, rendering this method more convenient for practical applications such as these.

To conclude, we believe that iDOP-PCR, employing the unique DNA polymerase properties and primer design, will become an important member of the WGA methods family. It provides simplicity, reproducibility and robustness in applications where fast and reliable amplification of genome copies are required.

## Acknowlegments

We thank Dr. Ya.I. Alekseev (All-Russia Institute of Agricultural Biotechnology, Moscow), Syntol JSC (Moscow, Russia), Evrogen JSC (Moscow, Russia) and Bioline Ltd (London, UK) for support of this project.

## Supporting information

**S1 Fig. Electrophoretic analysis of WGA libraries by agarose-gel electrophoresis.** The libraries were obtained by DOP-PCR, PicoPlex and iDOP-PCR methods from 15 pg (Lanes 1) and 0 pg (negative controls, Lanes 2) of the input human gDNA. **M** – 1 kb DNA Ladder.

**S1 Table. Average yield of DNA amplified by DOP-PCR, PicoPlex and iDOP-PCR.** Data for each point were obtained from analisis of 6 WGA samples.

**S2 Table. Multiplex STR genotyping of WGA samples and non-amplified gDNA.** In each wgaDNA and non-amplified gDNA sample, 38 allels were analyzed. Statistic data for each starting amount of gDNA amplified by each WGA method were obtained from the assay of six separate wgaDNA samples. Total N = 6 × 38 allels = 228 allels (100%). Allele concordance was calculated as a percentage of concordant alleles (Table A in S2 Table). Allele drop out (ADO) was calculated as a percentage of dropping-out concordant alleles (Table B in S2 Table). Allele drop in (ADI) was calculated as a percentage of dropping-in discordant alleles (Table C in S2 Table).

